# A DNA segregation module for synthetic cells

**DOI:** 10.1101/2022.04.30.489979

**Authors:** Mai P. Tran, Rakesh Chatterjee, Yannik Dreher, Julius Fichtler, Kevin Jahnke, Lennart Hilbert, Vasily Zaburdaev, Kerstin Göpfrich

## Abstract

The bottom-up construction of an artificial cell requires the realization of synthetic cell division. Significant progress has been made towards reliable compartment division, yet mechanisms to segregate the DNA-encoded informational content are still in their infancy. Herein, droplets of DNA Y-motifs are formed by liquid-liquid phase separation (LLPS). Entropy-driven DNA droplet segregation is obtained by cleaving the linking component between two populations of DNA Y-motifs. In addition to enzymatic cleavage, photolabile sites are introduced for spatio-temporally controlled DNA segregation in bulk as well as in cell-sized water-in-oil droplets and giant unilamellar lipid vesicles (GUVs). Notably, the segregation process is slower in confinement than in bulk. The ionic strength of the solution and the nucleobase sequences are employed to regulate the segregation dynamics. The experimental results are corroborated in a lattice-based theoretical model which mimics the interactions between the DNA Y-motif populations. Altogether, engineered DNA droplets, reconstituted in GUVs, could represent a strategy towards an entropy-driven DNA segregation module within bottom-up assembled synthetic cells.

**Table of Contents:** An entropy-driven DNA segregation module for bottom-up assembled synthetic cells is realized. It is based on DNA droplets that are engineered to segregate upon enzymatic or photocleavage inside giant unilamellar lipid vesicles (GUVs). The segregation kinetics is altered by the confinement, as confirmed by lattice-based numerical simulations. DNA segregation is further controlled by temperature, ionic strengths and nucleobase sequence.

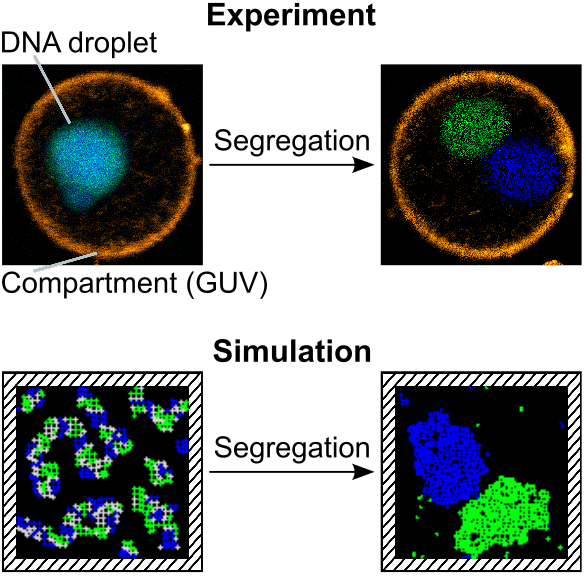

## 1 Introduction

A bottom-up assembled artificial cell should possess the ability to proliferate and evolve, which entails growth, division and information distribution.[1] The reconstitution of cell division has therefore gained growing interest.[1, 2, 3] A variety of strategies have been developed to divide lipid vesicles as synthetic cellular compartments.[4, 5, 6, 7, 8, 9] Some have also succeeded in encapsulating DNA or RNA inside dividing vesicles,[4, 10, 11] however, information was distributed by random partitioning only.

Thus, there is an undeniable call for a controlled DNA segregation module in artificial cellular systems.[1] The natural machinery for DNA segregation, namely the eukaryotic mitotic spindle or the bacterial Par system, are, for now, too intricate to be reconstituted and coordinated in a synthetic cell.[1, 12] Yet entropy-driven segregation of dense polymers in confinement, which is biologically relevant e.g. for bacterial chromosome segregation, could provide a feasible strategy.[1, 13, 14] Dense DNA packing could be achieved in synthetic cells by making use of a related physical mechanism, namely the process of liquid-liquid phase separation (LLPS). Several cellular organelles, most notably the nucleolus,[15] are manifestations of this phenomenon.[16] The benefits of passive localization and increased local concentration of chemical components due to LLPS have been exploited to create hierarchical subcompartments in synthetic cells.[17, 18, 19] Some of them have been engineered from DNA only, termed DNA droplets,[20, 21, 22, 23] capitalizing on advances in DNA nanotechnology.[24] The formation of DNA droplets by LLPS has been studied using coarse-grained modeling and analytical scaling theory.[25, 26] However, DNA droplet fission has not yet been simulated, although there is one example of a promising experimental mechanism.[22] Moreover, DNA droplets and their fission have not yet been reconstituted in confinement.

Here, we explore whether DNA droplet fission can be orchestrated in the confinement of water-in-oil droplets and giant unilamellar lipid vesicles (GUVs) to explore routes towards a DNA segregation module for synthetic cells. For this purpose, we report a system where DNA droplets can be segregated not only by enzymatic activity but also by light illumination with full spatiotemporal control. The segregation dynamics are characterized experimentally and recapitulated by lattice-based theoretical model in bulk and in confinement conditions. Reconstituted inside a GUV, we demonstrate that a fully synthetic DNA-based condensate could be a promising candidate for mimicking the nucleus of a synthetic cell.

## 2 Results and Discussion

### 2.1 Entropy-driven DNA segregation strategy

To build our synthetic DNA segregation module, we first need to assemble DNA coacervate droplets that are capable of undergoing fission. We adapted a design based on two types of DNA Y-motifs (**Figure 1a**, blue and green), whereby each Y-motif is formed from three single-strands of DNA which self-assemble into a Y-shaped motif with a planer end-to-end distance of approximately 10nm.[22] The Y-motifs possess two orthogonal types of palindromic eight nucleotide long sticky end sequences which allow for their polymerization among the same population (blue-blue, green-green). The two populations of Y-motifs are inter-connected by a six-arm linking motif, such that a single coacervate droplet forms (containing the blue and green Y-motifs as well as the linking motif, see Experimental Section). The cleavage of the linking motifs removes the interactions between the two DNA Y-motif populations, leading to their entropy-driven segregation into two separate liquid phases. Once the segregation process is completed, the daughter DNA droplets are spatially segregated due to the sequence-orthogonality of their interactions (Figure 1a).[22]

**Figure 1:**
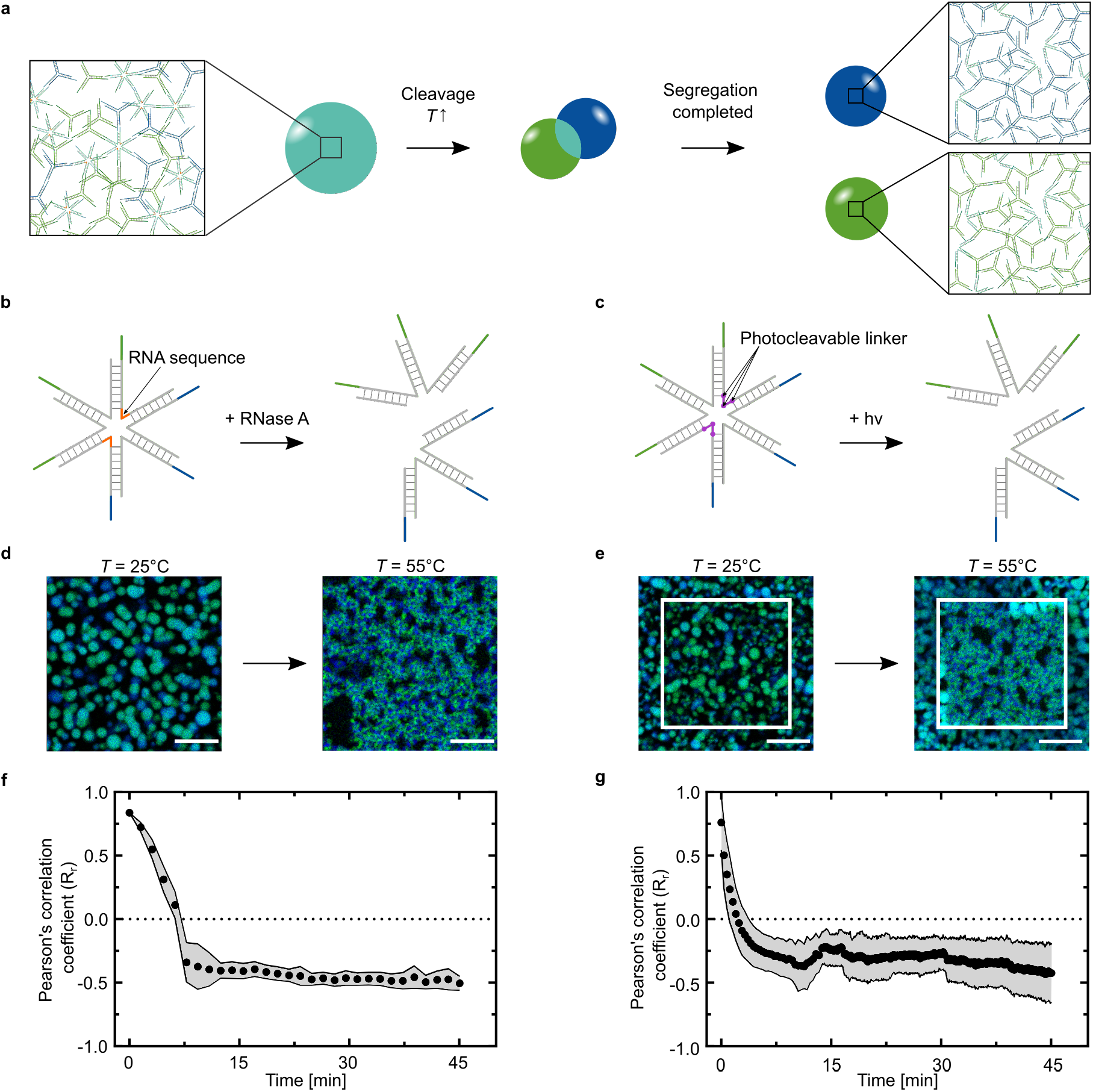
DNA segregation mechanism by enzymatic or photo-induced cleavage. a) Schematic of the DNA droplet consisting of two types of DNA Y-motifs (green and blue) with self-complementary sticky end sequences. A six-arm DNA motif interconnects the two populations to form a single droplet. Upon cleavage of the six-arm linking star and sufficient temperature increase, the initial droplet (cyan sphere) liquid-liquid phase separates into two segregated droplet populations (green and blue). b,c) Cleavage of the gray six-arm linking motif is achieved by enzymatic activity of ribonuclease A (RNase A) (b) or by light-triggered photocleavage of nitrobenzyle groups (c). d) Confocal overlay images of DNA droplets pre-treated with RNase A in bulk (*c* = 20 μgmL^-1^) before (left) and after heating to *T* = 55 °C for *t* = 4 min (right). e) Confocal overlay images of DNA droplets in bulk before (left) and after heating (right). The droplets within the region of interest (highlighted with a white square) were illuminated at *λ* = 405nm for *t* = 30 s to induce photocleavage. The two DNA populations were labeled with ATTO-488 (*λ_ex_* = 488nm) and ATTO-647N (*λ_ex_* = 640nm). Scale bars: 20 μm. f,g) DNA segregation dynamics of the RNase A-cleaved DNA droplets (f) and the photocleaved DNA droplets (g). Colocalization analysis (Pearson’s correlation) was performed and Pearson’s *R_r_* values are plotted over time (mean ± s.d, *n* = 16 regions in (f) and *n* > 64 regions in (g)).

For the cleavage of the linking motifs, we implemented two different strategies, namely, enzymatic cleavage and light-triggered cleavage (Figure 1b,c). Light-triggered cleavage allows for full spatiotemporal control over the segregation process, in particular within compartments. For enzymatic cleavage, ten DNA bases were replaced with RNA in two opposite strands at the centre of the six-arm linking DNA motif as proposed by Sato *et al*. (Table S1).[22] We can therefore make use of RNase A activity to degrade the RNA-DNA hybrid strands and hence the connection between the two halves of the six-arm motif.[22] In this way, the connection between the two populations of the Y-motifs is disrupted.

To enable photoinduced segregation, three photoreactive nitrobenzyle linkers were introduced within the same two strands of the six-arm motif that were modified for enzymatic cleavage (Figure 1c; Table S1, Supporting Information). The three photocleavable groups were positioned such that two five nucleotide long DNA oligos are released upon illumination and the six-arm motif splits into two.

We first tested these two DNA droplet segregation methods in bulk. After treatment with RNase A and heating to 55 °C to bring the DNA droplets to the liquid state, the DNA droplets undergo segregation in the entire observation chamber (Figure 1d). Despite activity of RNase A within a broad temperature range from 15-70 °C,[27] enzymatic activity cannot be spatially controlled in the bulk sample. Photocleavage, instead, could be triggered locally by illumination with the 405nm laser of the confocal microscope. DNA droplets in the illuminated area (highlighted by the white box in Figure 1e) undergo segregation, while adjacent droplets only fuse due to the heating.

To describe the segregation kinetics in a quantitative manner, we determined Pearson’s R_*r*_ correlation coefficient as a measure for the colocalization of the two Y-motif populations over time. Note that a value of R_*r*_ = +1 corresponds to perfect colocalization whereas R_*r*_ = −1 is obtained for perfectly anticorrelated data. The correlation coefficient decays and reaches a plateau within less than 10 min for both the enzymatic and the light-triggered DNA droplet segregation (Figure 1f,g). However, the decay kinetics differ for the two systems. Interestingly, we observed a latency for the enzymatic segregation (Figure 1f), while the photoinduced segregation follows an exponential decay (Figure 1g). This behaviour is consistent at different temperatures (Figure S1b-iii,iv, Supporting Information). Light can penetrate across the entire droplet and cleave all linking DNA motifs simultaneously, while enzymatic activity is initially limited to the surface of DNA droplets in a gel state due to the size exclusion.[28] Once the DNA droplet is in a liquid state, enzymes can have access to the rest of the droplet. This ongoing enzymatic reaction delays the segregation process, explaining the delayed decay of the Pearson’s correlation coefficient for the enzymatic cleavage.

We have thus demonstrated enzymatic and photoinduced cleavage as two distinct mechanisms for the entropy-driven segregation of densely packed droplets of DNA Y-motifs, whereby photo-cleavage provides full spatiotemporal control. We will therefore use the light-triggered segregation for the following experiments.

### 2.2 DNA segregation in cell-sized confinement

Having demonstrated DNA segregation in bulk, we next introduced geometric confinement by encapsulating the DNA droplets into cell-sized compartments. For this purpose, we encapsulated the DNA droplets in water-in-oil droplets and carried out the experiment analogous to the bulk segregation. Importantly, we find that the confinement slows down the segregation kinetics (**Figure 2**a; Video S1, Video S2, Supporting Information). Tracing the colocalization of the two populations over time shows full segregation in bulk after 10 min while the segregation process is still not fully completed after 45 min in confinement (see graph in Figure 2a). This result is consistent for both enzyme- and photo-induced segregation (Figure 2a, Figure 1f; Figure S1a,b-iii, Supporting Information).

**Figure 2:**
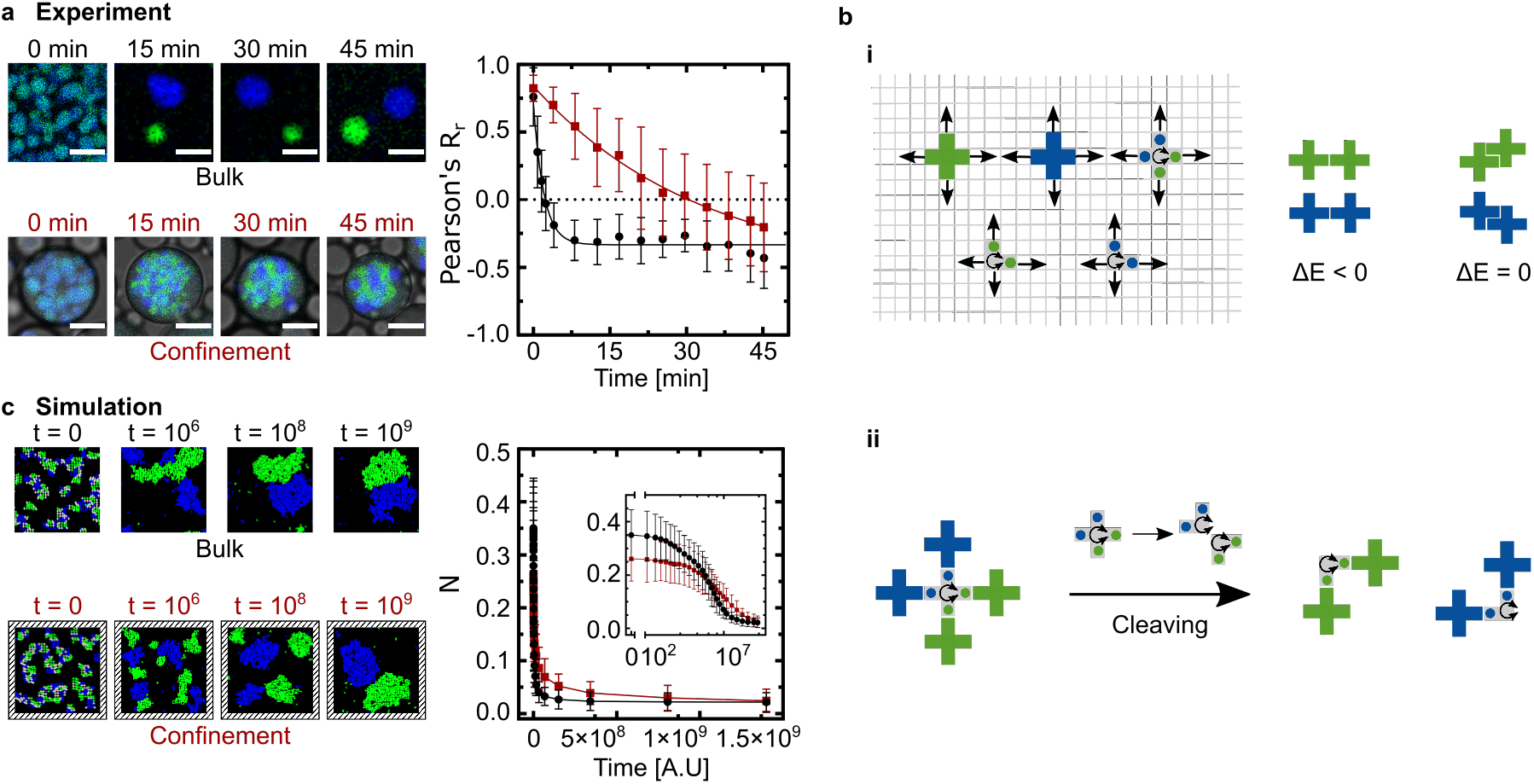
Effects of geometrical confinement on DNA segregation dynamics. a) Experimental DNA segregation dynamics in bulk compared to in confinement. Left panel: Representative confocal microscopy images of different time points showing DNA droplet segregation in bulk (upper, fluorescence overlay) and in water-in-oil droplets (lower, fluorescence and brightfield overlay). The two DNA populations were labeled with ATTO-488 (*λ_ex_* = 488nm) and ATTO-647N (*λ_ex_* = 640nm). Scale bars: 10 μm. Right panel: Pearson’s *R_r_* values plotted over time for DNA segregation process in bulk (mean ± s.d, *n* > 64 regions, black circles) and in confinement (mean ± s.d, *n* > 70water-in-oil droplets, red squares). The data is fitted using one-phase decay equation (shown in lines). b) Schematic of implemented lattice-based model. i) Cross-shaped DNA-motifs can perform next nearest neighbor diffusion in four directions, linking (gray) and cleaved “L”-shaped motifs can additionally undergo stochastic rotations. The system free energy is only reduced for arm-to-arm configurations and not affected by any other contact. ii) Cleavage mechanism in the model: A mixed phase is produced from a random initial configuration, and thereafter cleaving of linking motifs give rise to two “L”-shaped motifs. c) Simulated DNA segregation dynamics in bulk compared to that in confinement. Left panel: Snapshots of DNA droplets undergoing segregation with periodic boundary conditions, mimicking experimental bulk condition (upper) and within rigid boundary, mimicking experimental confinement (lower). The simulation time step (t) is indicated above each snapshot. For visibility, the gray particles are colored in the same color as their population (see Figure S2, Supporting Information for original images). Right panel: Average number of nearest neighbors of the other DNA population over time in periodic boundary (mean ± s.d, *n* > 10^3^ (ensemble average), black circles and line) and in rigid boundary (mean ± s.d, *n* > 10^3^ (ensemble average), red squares and line).

To shed light on the possible mechanism and to confirm that the geometric confinement indeed can slow down segregation of DNA droplets, we considered a lattice-based model of this process. The model describes each DNA Y-motif as a cross-shaped particle diffusing on a two-dimensional (2D) square lattice in the four possible directions (up-down-left-right) with defined particle-particle interaction strengths and excluded volume effects. Systems of cross-like particles on a lattice are a well-studied model of statistical physics.[29, 30, 31, 32] At the same time, this model contains the minimal geometry necessary to account for all inter- and intra-species interactions as happening in experiments. Therefore, whilst being simpler than a hexagonal lattice model, the square lattice still captures the experimental system. Analogous to the experiment, the model contains three types of particles, namely two populations which interact amongst themselves (blue and green) and linking particles (analogous to the six-arm DNA motif) which interconnect the two populations. When the linking particles are cleaved, they produce two “L”-shaped particles of the respective DNA populations (Figure 2b). Both the linking motifs and the cleaved “L”-shaped motifs can rotate stochastically to maintain the orientational uniformity of the system. The overall surface coverage by the particles is kept below 0.8 to ensure that the system maintains a liquid state.[33]

We reproduced the bulk and confinement conditions by using periodic and rigid boundaries, respectively. Starting from a random initial configuration with a given particle density, interaction strengths and system temperature, a mixed phase is produced to mimic the initial mixed DNA droplets as observed in the experiment (Figure 2c). After cleaving the linking motifs, a higher temperature is applied to the mixed phase so that the linking motifs break apart to produce two “L”-shaped motifs, each belonging to their respective DNA population. Similar to the experiment, we observed phase segregation resulting in two separate droplet species. For the quantitative analysis of the simulations, we introduced a parameter *N*, which defines, for a single blue or green DNA motif, the average number of nearest neighbor sites occupied by the motifs of the other population. This metric is qualitatively equivalent to the Pearson’s correlation coefficient *R_r_* measured in the experiment, whereby higher number of different motif neighbours corresponds to a mixed state while lower number is indicative of segregated droplets. Importantly, in agreement with experimental results, the phase segregation in confinement is slower. The slower segregation kinetics in confinement are likely due to particles which detach in close proximity to the compartment boundary, which attach back in close proximity to their initial position. This slows down the overall reorganization of the DNA droplets. This finding corroborates the role of cell-sized confinement on our DNA segregation process, accentuating the importance of including compartments while studying and constructing cellular components.

### 2.3 Control of DNA segregation kinetics

Having demonstrated that DNA segregation is slowed down in confinement, we set out to elucidate mechanisms by which DNA segregation can be sped up and controlled in a compartment. We therefore probed different strategies to tune the interaction strength and the diffusion of the DNA Y-motifs. The most obvious parameter which will influence both, DNA-DNA interaction and diffusion, is temperature. We examined the segregation process at 35-65 °C which covers the gel to liquid transition of DNA droplets.[22] At 35 °C and 45 °C, the cleaved droplets remain unchanged in both enzymatic and photocleavable systems (**Figure 3**a-i,ii; Figure S1b-i,ii, Supporting Information). Neither fusion nor fission events occurred, indicating that the DNA droplets are in a gel state. When the temperature was raised to 55 °C after incubation at 35°C or 45 °C, DNA segregation was observed (Figure S3, Supporting Information). A temperature increase from 55 °C to 65 °C significantly speeds up the DNA segregation (Figure 3a-iii,iv; Figure S4, Figure S5, Supporting Information). Complete segregation is detected in nearly all water-in-oil droplets after only 5 min of incubation at 65 °C (Figure 3a-iv), whereas the process was not fully completed at 55 °C even after 45 min (Figure S6, Supporting Information). At 65 °C, we additionally observed partial wetting of the blue DNA population on the water-in-oil interface (Figure S6, Supporting Information). Wetting is enabled by the presence of negatively charged Krytox molecules in the surfactant layer.[34, 35] It is interesting that it is only observed for the blue population. This is likely due to its lower unpaired fraction at elevated temperatures as revealed by a comparative analysis of the melting profiles of the two sequences (Figure S7, Supporting Information).

**Figure 3:**
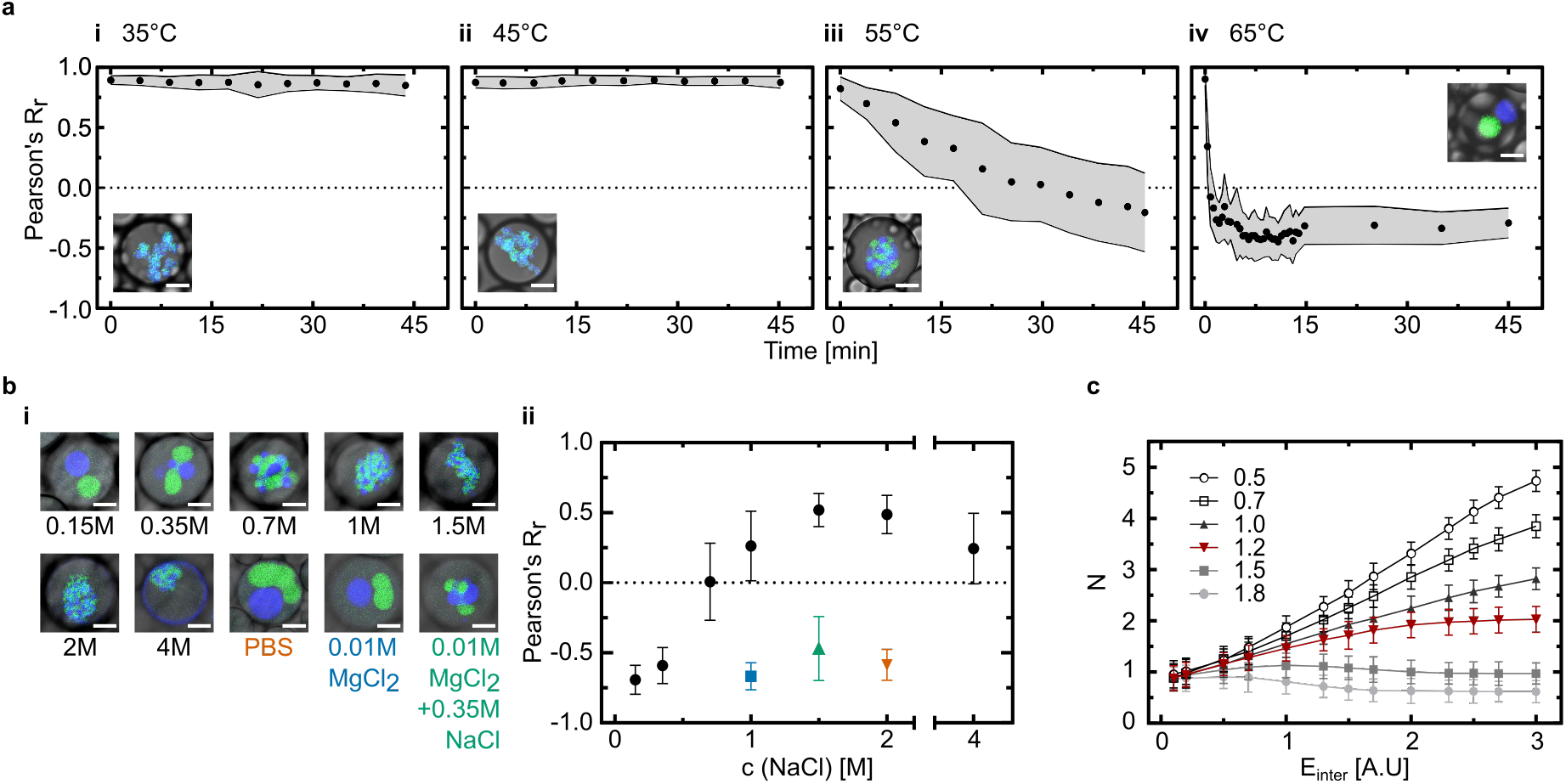
DNA segregation dynamics depends on different parameters. a) Temperature dependence. Photo-induced segregation process is observed over time in different water-in-oil droplets at different temperatures (i-iv) (*t* = 45 min, *T* = 35-65 °C, 10 °C step) and Pearson’s *R_r_* values are plotted (mean ± s.d, *n* > 60water-in-oil droplets). Graph insets: overlay of confocal fluoresecnce and brightfield images of representative water-in-oil droplets in each condition. The fluorescent pixels outside of the DNA droplets were removed for visibility. b) Ionic strength dependence. i) Overlay of confocal fluorescence and brightfield images of photo-cleaved DNA droplets in water-in-oil droplets after heat treatment (*t* = 10 min, *T* = 55 °C) at different ionic conditions. ii) Pearson’s *R_r_* values of the two DNA populations plotted against NaCl concentrations (black circles) and other ionic conditions in (i) (PBS in orange, 0.01M MgCl_2_ in blue and 0.01M MgCl_2_ + 0.35M NaCl in green, mean ± s.d, *n* > 70water-in-oil droplets). Scale bars: 10 μm c) Lattice simulations: Average number of neighbors *N* as a function of inter and intra-population strengths for several values of the linear multiplicative constant 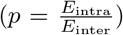 ranging from 0.5 to 1.8. At the value *p* = 1.2, the curve (red line) is qualitatively similar to the dependence in experiment. Now, for *p* <1.2, the intra-population interactions dominate and *N* increases as **E**_inter_, leaving the system in a more mixed state, whereas for *p* > 1.2, the system is largely segregated into two DNA populations.

Ionic strength, on the other hand, changes the DNA interaction strength while having only a minor influence on the diffusion rates. We thus formed the DNA droplets as previously in 0.35M NaCl, 1× Tris-EDTA pH 8.0, pelleted and re-suspended the droplets in solutions of different ionic strengths prior to encapsulation into water-in-oil droplets (see Experimental Section, Figure 3b). We then imaged the droplets after photocleavage and 10 min incubation at 55 °C. As the concentrations of NaCl increases from 0.15M to 1.5M, the correlation coefficient increases from −0.69 to 0.52, respectively, indicating reduced DNA segregation (Figure 3b). At 2M and 4M NaCl, we observed wetting interactions between the blue Y-motifs and the surfactant-stabilized water-oil interface (Figure 3b-i). At high concentrations of monovalent salt, charge screening is effectively leading to an interaction between the DNA and the negatively charged Krytox in the surfactant layer.[34] 10mM of divalent cations result in similar segregation kinetics to 0.15M NaCl. Due to the more effective charge screening of divalent cations, more than 10 times lower concentrations are required here.

We hypothesize that increasing ionic strength increases the interactions between Y-motifs of the same type as well as the unspecific electrostatic interactions of all Y-motifs. To test this hypothesis, we simulated the DNA segregation process for different inter- and intra-population attractive interaction strengths. In absence of any known explicit dependence among different interaction strengths in the experiment, we here assume a linear relation with a multiplicative constant 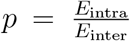, where *E*_intra_ and *E*_inter_ represent the attractive interaction among similar (intra-population, i.e. green-green, blue-blue) and different (inter-population, i.e. green-blue) types of DNA-motifs, respectively. In the experiment, intra-population interactions are mediated by base-pairing and electrostatic interactions; inter-population interaction is mediated by electrostatics only. The simulation reveals that, depending on the values of p, the behaviour of the segregation process responds differently to varying interaction strengths, as shown in Figure 3c. As p increases, DNA motifs of the same population form relatively robust clusters, reflected in a reduction in *N*, whereas decreasing p has the opposite effect (Figure 3c). At p=1.2, the curve has qualitatively the same shape as in the experiment. This indicates a possible approximate relation between the two types of interactions (*E*_intra_ and *E*_inter_), whereby the interaction between the same population is enhanced by a factor of 1.2 more strongly at increased ionic strength. This appears reasonable as both base-pairing as well as electrostatic interactions are enhanced at increased ionic strength. Intra-population interactions, which are mediated by base-pairing and electrostatic interactions, should thus be affected more strongly. We can deduce that electrostatics contribute by about 20% to the overall interaction, where base-pairing interactions dominate with 80 %.

### 2.4 Towards a DNA segregation module for a bottom-up assembled synthetic cell

The evolvability of living systems requires a correlation between genetically encoded information (genotype) and function (phenotype). It is thus interesting to explore if the DNA segregation phenotype can, in principle, be controlled by the DNA sequence. The sequence of the sticky ends determines the strength of the intra-population interactions without affecting the interactions between the two different types of Y-motifs. According to our previous simulation results (Figure 3c), we thus expect a sequence dependence of the segregation behaviour. More interestingly, the sticky end sequence can also impact the mechanical properties of the link, e.g. if unpaired bases are introduced which enhance the flexibility. We thus introduced one single free base between the arm of the Y-motif and its sticky end as illustrated in **Figure 4**a. We observed a noticeable effect on the droplet formation for the blue Y-motif population (Figure S8, Supporting Information), in agreement with previous reports.[25] Despite the higher melting temperature of the blue Y-motif compared to the green population (Figure S7, Supporting Information), this Y-motif forms more unstable droplets than the other population independent of the choice of fluorophores (Figure S9, Supporting Information).[36] This is likely due to the lower fraction of unpaired bases of this population at elevated temperatures as revealed by an analysis of the melting profile (Figure S7, Supporting Information). It is therefore interesting to study the effect of flexibility on the segregation process.

**Figure 4:**
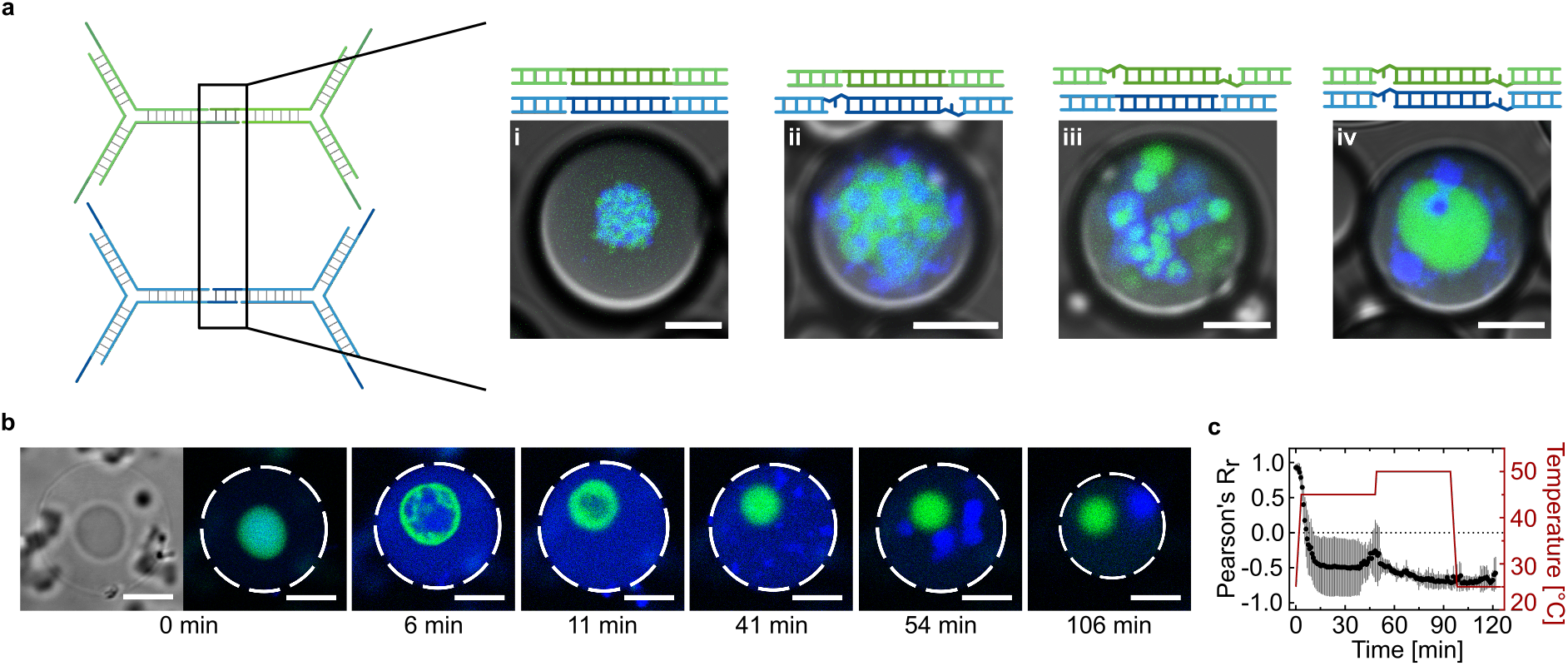
A DNA segregation module for synthetic cells. a) DNA sequence controls segregation phenotype. Confocal overlay images of photo-cleaved DNA droplets in water-in-oil droplets after heat treatment (*t* = 5 min, *T* = 50 °C). New sequences were designed by inserting one free base at the 3’ end of both types of sticky ends (green and blue) and four combinations were generated from these four types of sticky ends (i-iv). The two DNA populations were labeled with ATTO-488 (*λ_ex_* = 488nm) and ATTO-647N (*λ_ex_* = 640nm). b) Confocal time series of a single DNA droplet inside GUV (brightfield image on the left) fully segregated into two daughter DNA droplets upon illumination (*λ* = 405nm, *t* = 30 s, *I* = 5mW) and heat application. The two DNA populations were labeled with ATTO-488 (*λ_ex_* = 488nm) and ATTO-647N (*λ_ex_* = 640nm). Scale bars: 10 μm. c) Colocalization analysis of the two DNA populations (black circles, mean ± s.d, *n* = 17 z-planes) and the temperature profile (red line) over time.

We find that increased flexibility alters not only the DNA droplet formation (Figure S10, Supporting Information) but also the DNA segregation. Upon cleavage at room temperature, we find that aggregations of modified populations have formed outside the droplet (Figure S11, Supporting Information), whereas DNA segregation required temperatures of at least 55 °C in previous experiments. Remarkably, the addition of the free adenine changed the spatial trajectory of the segregation process. With the original less flexible design, the DNA droplets undergo spinodal decomposition where the two populations become patchy before full segregation. Now, in the cases where only one type of Y-motifs is modified, we observed a hierarchical core-shell organization within the DNA droplets where the modified population is the core shielded by the other population (Figure 4a-ii,iii). The addition of the free base to both populations restored the spinodal decomposition, however, with a more separated product (Figure 4a-iv). These changes in spatial organization imply a change in the surface tension of the droplets which might be due to the exposed hydrophobic base of the unpaired adenine, which effectively acts as a surfactant. RNA, which displays secondary structures exposing its bases, are a prominent constituent of many cellular condensates.[37] This poses questions on how much secondary structures and DNA unwinding influence the behaviours of their condensates and whether the cell also uses this to regulate its functions. In summary, we have shown that, with the DNA sequence, we can not only regulate the kinetics of the DNA segregation but also the segregation trajectory in confinement.

Beyond sequence programmability, another important feature of a DNA segregation module for synthetic cells is that it should ultimately function inside of lipid vesicles. As the next step, we thus reconstituted the DNA droplets in giant unilamellar lipid vesicles (GUVs). Since the reconsititution in water-in-oil droplets was successful, we decided to use the droplet-stabilized GUV formation method, which uses water-in-oil droplets as a template for the formation of free-standing GUVs.[35, 34] In brief, we encapsulated all components required for the DNA segregation together with small unilamellar lipid vesicles (SUVs) in water-in-oil droplets. The SUVs fuse at the water-oil-interface to form a spherical supported lipid bilayer, which can be released from the droplet as a conventional GUV. After GUV formation, the single-stranded DNA is annealed to form the initial DNA droplet in GUVs. The segregation process was successfully triggered by RNase A activity (Figure S12, Supporting Information) or locally by light (Figure 4b; Figure S13, Video S3, Video S4, Supporting Information). The light-driven segregation is captured over time and colocalization was analyzed for a quantitative view of the segregation process (Figure 4b,c). By reproducing DNA segregation in GUVs, we come closer to a controllable analogue of cell division which would open new doors for the idea of constructing a synthetic cell from scratch.

## 3 Conclusion

The segregation of informational molecules like DNA, is a hallmark of natural cell cycle, which makes the quest for a mechanism of DNA spatial segregation pivotal for the construction of a synthetic cell. Bottom-up synthetic biology has recently witnessed a myriad of successful and reliable approaches to divide lipid vesicles as synthetic cellular compartments. Now, it will be important to combine compartment division with suitable mechanisms for DNA segregation. Random partitioning, entropy-driven segregation and the reconstitution of the biological machinery have been proposed for this purpose.[1] Here, we realize an entropy-driven DNA segregation module in water-in-oil droplets and GUVs, which makes use of the physical principles of liquid-liquid phase separation. It provides more control than random partitioning, while maintaining a relatively simple set of components. It is conceivable that early synthetic cells could function with a membrane-free nucleus mimic. However, the downside of our system is the fact that at least the sticky end sequences of the daughter droplets cannot be identical. Therefore, the realization of multiple growth and division cycles would require a mechanism for regeneration. Yet, studying DNA droplets and their segregation kinetics is interesting in itself, as it provides a model system to shed light on the relation between DNA packing and access of DNA binding proteins like transcription factors. We have seen an example of this when revealing the different segregation kinetics for the enzymatic and the photo-induced segregation. Notably, the role of geometric confinement was observed to play a role in slowing down the segregation process, emphasizing compartmentalization as a requisite for the apprehension of cellular systems. Furthermore, a lattice-based model could capture the phenotypic behaviors of DNA droplet segregation. We also demonstrate orchestration of the system dynamics via temperature, ionic strengths and nucleobase sequences. With such versatile control toolbox, our system is not limited to the biomimicry of biocondensates but can be used as an engineerable model system for studying physical properties of LLPS and nucleotide interactions of natural cells. Our further implementation of the process into GUVs does not only bring the system closer to its natural analogue but also allows, in principle, for feeding mechanisms using targeted vesicle fusion to be applied for recurrent segregation.[38, 6] As a next step, it will be crucial to combine the DNA segregation modules with existing modules for compartment division.[6] Altogether, our work presents a physical approach towards providing a functional DNA segregation module for synthetic cells. Furthermore, our results will not only benefit the development of a fully synthetic cell but also prompt new questions on the role and properties of DNA in subcellular condensates.

## 4 Experimental Section

### 4.1 Annealing of DNA droplets

The oligonucleotides were purchased from Integrated DNA Technologies (IA, USA, purification: cartridge (OPC)–grade purification); modified oligonucleotides were purchased from biomers.net (Ulm, Germany, purification: high-performance liquid chromatography (HPLC)). The oligonucleotides were dissolved at 100 μM in 10mM Tris-HCl buffer (pH 8.0) containing 1mM EDTA purchased (Merck Millipore, MA, USA) and stored at −20 °C for further use.

The DNA strands were mixed in an Eppendorf PCR tube at different concentrations as described in Table S2 (Supporting Information) in 10mM Tris-HCl buffer (pH 8.0) containing 1mM EDTA and 350mM NaCl. The DNA strands were then annealed on a C1000 Touch™ Thermal Cycler from Bio-Rad (CA, USA). The sample was heated at 85 °C for 1 min and then cooled down to 25 °C at a rate of −1 °Cmin^-1^. The annealed DNA droplets were then pelleted using a Centrifuge 5804 from Eppendorf (Hamburg, Germany) at 10 000 g at 4 °C for 15 min. The supernatant was then discarded and the DNA droplets were re-suspended in the buffer of choice depending on the experiment (see Results and Discussion). The volume added for resuspension is 0.25× the initial volume. The concentrated DNA droplets were then stored for up to three days at room temperature in a dark box to prevent photocleavage or used immediately for further experiments.

### 4.2 Enzymatic degradation of DNA-RNA strands

RNase A purchased from Promega (WI, USA) was dissolved in deionized water and added to the concentrated DNA droplet solution at a final concentration of 20 μg mL^-1^. The samples were then imaged after a minimum incubation of 15 min in bulk or in water-in-oil droplets.

### 4.3 Light-triggered DNA cleavage

The sample containing DNA droplets was put onto the observation chamber. The light-triggered cleavage was either achieved by incubation for 5 min under a UV lamp (Hamamatsu) equipped with a 365nm filter or local illumination using the laser of the confocal laser scanning microscope at a wavelength of 405nm and a power of 5mW for at least 30 s.

### 4.4 Preparation of the heated observation chamber

Standard range smart substrates (SmS) measuring 18mm by 18mm with a thickness of 0.17mm were purchased from Interherence GmbH (Erlangen, Germany). For experiments with GUVs, the SmS was coated with 1% (w/v) BSA (bovine serum albumin, SERVA Electrophoresis GmbH, Germany) for at least 5 min to prevent fusion of the GUVs with the glass surface. After the BSA coating, the SmS was washed with deionized water and dried under an airflow. A coverslip (measuring 9mm by 6mm with a thickness of 0.17mm) was assembled onto the heating region of the SmS using double-sided sticky tape. The sample solutions were immersed into the slit between the SmS and the coverslip, and the edges were sealed with two component dental glue.

### 4.5 Confocal microscopy

The samples in the custom-built observation chamber were imaged with a confocal laser scanning microscope LSM 900 (Carl Zeiss AG, Oberkochen, Germany). The pinhole aperture was set to one Airy Unit and heat required experiments were performed with the temperature control unit VAHEAT from Interherence GmbH (Erlangen, Germany). The images were acquired using a 20× objective (Plan-Apochromat 20×/0.8 M27, Carl Zeiss AG). Images were analysed and processed with ImageJ (NIH, brightness and contrast adjusted). For images in Figure 2a, background pixels were removed using Remove Outliers function of ImageJ (radius: 2, threshold: 10).

### 4.6 Image processing

All microscope images were processed using ImageJ (NIH). For DNA segregation in bulk, an image was split into 16 equal regions and processed individually. For DNA segregation in water-in-oil droplets, each water-in-oil droplet was processed individually. The water and oil interface was defined manually and all pixels outside water-in-oil droplet were removed. Except for experiments in water-in-oil droplets, if a z-stack of the sample was imaged, each slice was processed separately. In the case of DNA segregation in GUVs, pixels from region outside the GUV and artefacts (signal from DNA droplets in solution outside of GUV) were removed. A Gaussian blur with a sigma of 2 was then applied on each region of interest. Images were then saved in TIFF files for further analysis.

### 4.7 Data analysis

The Pearson correlation coefficient *R_r_* was calculated using the formula:

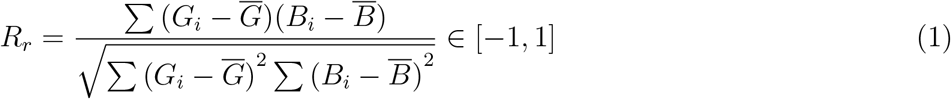

where *G_i_* or *B_i_* is the intensity of the i^th^ pixel in the green or blue channel respectively; 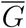 and 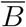 are the mean values of all pixel intensities of each channel. Images were analyzed using MATLAB ver. R2018b (code provided in Supporting data). Plot graphing and statistical tests were performed using GraphPad Prism.

### 4.8 Encapsulation of DNA droplets into water-in-oil droplets

For the formation of water-in-oil droplets, the DNA droplet-containing aqueous phase was layered on top of the oil phase in a volumetric ratio of 1:3 within a microtube (Eppendorf). For the segregation of DNA droplets using enzymatic activity, RNase A purchased from Promega (WI, USA) was added at a final concentration of 20 μg mL^-1^ before putting the aqueous phase onto the oil phase. Droplet formation was induced by manual flicking for about 10 times to produce cell-sized droplets. To stabilize the water-in-oil droplets, perflouro-polyether-polyethylene glycol (PFPE-PEG) block-copolymer fluorosurfactants (008-PEG-based fluorosurfactant) purchased from Ran Biotechnologies, Inc. (MA, USA) was dissolved in HFE-7500 oil purchased from DuPont (DE, USA) at a concentration of 3 % v/v to withstand high temperature.

### 4.9 Lattice-based model and numerical simulation

The DNA segregation model is revisited by numerical simulation in both bulk and confinement conditions. The bulk condition in simulation is realized by a periodic boundary, where the DNA motifs can exit from one side of the boundary and re-enter from the other side vertically and horizontally. Whereas in the confinement condition, the DNA motifs can not pass through the boundary to re-enter, rather, they get obstructed by the rigid boundary and can only get reflected back to the system. The dynamic Monte Carlo simulation is run on a lattice system of dimension 100×100. The dynamics starts from a random initial configuration with same density of 0.019 for the three types of DNA populations: blue, green (two DNA motifs) and gray (linker DNA motif). A single Monte Carlo step (MCS) is defined as: when on average each DNA motif is attempted to move to any of the four directions with equal probability as a diffusion process and attempted to rotate in either direction clockwise or anti-clockwise with equal probability as a rotational diffusion process. A single run of the simulations contains maximum of 10^9^ MCS. The significant effect of rotation in terms of altering interaction energy only applies for gray linking DNA motifs before cleaving and “L”-shaped motifs after cleaving, apparently, the green and blue DNA motifs do not rotate. Movements and rotations take place using standard Metropolis algorithm, which involves change in interaction energy ΔE and the explicit temperature *T* of the system. The interaction strengths (*E*_intra_ and *E*_inter_) are defined with respect to *k_B_T* and the Boltzmann constant (*k_B_*) is taken as unity. As *T* increases, the system is more likely to be randomized by thermal moves. Before cleaving, we took *T* = 0.55, and after the cleaving, we increased it to 0.70, whereas the attractive interactions *E*_inter_ and *E*_intra_ were varied in the range of 0.2 to 3.0 to observe various regimes of the segregation process. To quantify the segregation process, we used the average number of neighboring sites (*N*) of the other population for a single DNA motif as a function of simulation time for fixed parameters such as interaction strengths and temperature as shown in Figure 2c. However, to view the dependence of *N* on interaction strengths, we measured *N* as an ensemble average over 10^3^ simulations with different initial configurations with each run for 10^9^ MCS as displayed in Figure 3c. Now, since a single cross DNA motif has eight nearest neighbors, *N* can reach the maximum value of 8 if all the neighboring sites are occupied by the other population. When *E*_inter_ is high, similar DNA motifs are more prone to form rigid cluster, subsequently decreases the value of N. In contrast to that when *E*_inter_ is low, the system appears to be in a mixed state of two DNA population, giving rise to higher number of N. For a specific value of 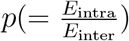, the simulation qualitatively recapitulates the experimental observations (Figure 3c). The numerical simulations were conducted using the code provided. We used Intel C-compiler to process the code.

### 4.10 SUV formation

1,2-dioleoyl-sn-glycero-3-phosphoglycerol (DOPG), and 1,2-dioleoyl-sn-glycero-3-phosphocholine (DOPC), were purchased from Sigma-Aldrich (Darmstadt, Germany), stored in CHCl_3_ at −20 °C and used without further purification. Small unilamellar vesicles (SUVs) were formed by mixing DOPG and DOPC at a 30%:70% molar ratio in a glass vial and later dried gently under a stream of nitrogen gas. The vial was kept under vacuum in a desiccator for 30 min to remove traces of CHCl_3_. The dried lipid film was resuspended in 10mM Tris-HCl buffer (pH 8.0) containing 1mM EDTA at a lipid concentration of 8mM. The solution was vortexed for at least 10 min to trigger liposome formation and subsequently extruded to form homogeneous SUVs with ten passages through a polycarbonate filter with a pore size of 100nm purchased from Avanti Polar Lipids, Inc.(AL, USA). SUVs were afterwards stored at 4°C for up to 3 days or used immediately for GUV formation.

### 4.11 GUV formation

To prepare the aqueous solution, the DNA droplet pellet was re-suspended in the initial buffer at a 6.67× increased concentration. The re-suspended DNA droplet solution was then denatured at 95 °C for at least 5 min on a C1000 Touch™ Thermal Cycler from Bio-Rad (CA, USA). The aqueous solution consisted of 25% v/v SUV solution (containing 8mM lipids as prepared above) and 60% v/v denatured DNA solution in 10mM Tris-HCl buffer (pH 8.0) containing 1mM EDTA and 350mM NaCl. The oil–surfactant mix contained HFE-7500 fluorinated oil (3M, Germany) with 1.4 wt% 008-PEG-based fluorosurfactant (RAN Biotechnologies, MA, USA) and 10mM PFPE–carboxylic acid (Krytox, MW, 7000-7500 gM^-1^, DuPont, Germany). The aqueous solution was layered on top of the oil-surfactant mix in a volumetric ratio of 1:3 inside a microtube (Eppendorf). The tube was manually shaken for about 30 s until water-in-oil droplets formed. The SUVs fused to form a spherical supported lipid bilayer at the droplet periphery and created droplet-stabilized GUV (dsGUV). The dsGUVs tube was stored at 4 °C for 3 h before release. The same buffer at equal volume to the aqueous solution was pipetted on top of the droplet layer. To destabilize the droplets for release, perfluoro-1-octanol (PFO) destabilizing agent (Sigma-Aldrich, Germany) of the same volume was added slowly. Within a minute, the milky emulsion disappeared and formed a transparent aqueous layer on top of the oil–surfactant mix. The released GUVs in the transparent aqueous layer were carefully removed with a pipette and transferred into another microtube. The released GUVs were stored at room temperature in a dark box for up to three days or used immediately for further experiments.

### 4.12 Formation of DNA droplets in GUV

To form DNA droplets inside the GUVs, the released GUVs were incubated on a C1000 Touch™ Thermal Cycler from Bio-Rad (CA, USA) at 85 °C for 1 min and then cooled down to 25 °C at a rate of −1 °Cmin^-1^. The denatured DNA inside the GUVs and in the solution re-annealed into DNA droplets. The solution was then either immediately put on an observation chamber for imaging or stored at room temperature in a dark box for up to three days before imaging.

## Acknowledgements

M.P.T and K.G received funding by the Federal Ministry of Education and Research (BMBF) and the Ministry of Science Baden-Württemberg within the framework of the Excellence Strategy of the Federal and State Governments of Germany. K.G. received funding from the Deutsche Forschungsgemeinschaft (DFG, German Research Foundation) under Germany’s Excellence Strategy via the Excellence Cluster 3D Matter Made to Order (EXC-2082/1 - 390761711) and the Max Planck Society. K.J. thanks the Carl Zeiss Foundation and Joachim Herz Foundation for financial support. R.C. and V.Z. were supported by “Life?” initiative of the Volkswagen Stiftung. L.H. is supported by the Helmholtz program Natural, Artificial, and Cognitive Information Processing (NACIP). The Max Planck Society is acknowledged for its general support. All authors thank Yusong Ye and Tim Klingberg for helpful discussions.

