## Supplementary Information for "A DNA segregation module for synthetic cells"

### Supporting information for:
### A DNA segregation module for synthetic cells

*Mai P. Tran  Rakesh Chatterjee  Yannik Dreher  Julius Fichtler  Kevin Jahnke  Lennart Hilbert  Vasily Zaburdaev*  Kerstin Göpfrich**

Mai P. Tran
Address: Biophysical Engineering Group, Max Planck Institute for Medical Research, Jahnstr. 29, 69120 Heidelberg, Germany
Department of Biosciences, Heidelberg University, 69120 Heidelberg, Germany

Dr. Rakesh Chatterjee, Prof. Dr. Vasily Zaburdaev*
Address: Friedrich-Alexander-Universität Erlangen-Nürnberg, Department of Biology, Cauerstraße 11, 91058 Erlangen, Germany
Max-Planck-Zentrum für Physik und Medizin, 91058 Erlangen, Germany

Julius Fichtler
Address: Biophysical Engineering Group, Max Planck Institute for Medical Research, Jahnstr. 29, 69120 Heidelberg, Germany

Yannik Dreher, Kevin Jahnke, Dr. Kerstin Göpfrich*
Address: Biophysical Engineering Group, Max Planck Institute for Medical Research, Jahnstr. 29, 69120 Heidelberg, Germany
Department of Physics and Astronomy, Heidelberg University, 69120 Heidelberg, Germany

Prof. Dr. Lennart Hilbert
Address: Institute of Biological and Chemical Systems, Karlsruhe Institute of Technology Hermann-von-Helmholtz-Platz 1, 76344 Eggenstein-Leopoldshafen, Germany
Zoological Institute, Department of Systems Biology / Bioinformatics, Karlsruhe Institute of Technology Fritz-Haber-Weg 4, 76131 Karlsruhe , 76134 Karlsruhe

### 1 Experimental section

#### 1.1 Annealing of DNA droplets without linking motifs

The DNA strands were mixed in an Eppendorf PCR tube at 5 µM per strand in 10 mM Tris-HCl buffer (pH 8.0) containing 1 mM EDTA and 350 mM NaCl. For the original droplets, G and B strands were used (see Table S1, Table S2). For the modified droplets, G-free and B-free strands were used. For labeling, 30% of G-2 and G-free-2 strands were replaced with G-2-FAM and 30% of B-2 and B-free-2 were replaced with B-2-DY405. The DNA strands were then annealed as described in Experimental section.

#### 1.2 Encapsulation of DNA droplets in fluorescently labelled GUV

SUVs were prepared using 1,2-dioleoyl-sn-glycero-3-phospho ethanolamine-N-(lissamine rhodamine B sulfonyl) (ammonium salt) (Liss Rhod PE, purchased from Sigma-Aldrich, Germany), DOPG, and DOPC mixed at 1%:30%:69% molar ratio. SUV formation was done as described in Experimental section.
The aqueous solution was then prepared and used for GUV formation using method described in Experimental section. For enzyme-induced DNA droplet segregation in GUV, DNA droplet pellet was resuspended in the initial buffer for a concentration of 4× then mixed with RNase A at a final concentration of $20\,\mu g\,mL^{-1}$.

#### 2 Table S1: Oligonucleotide sequences

Table S1: Oligonucleotide sequences.
The sequences are adapted from [1]. "G", "B" and "L" in the "Name" column are abbreviations for green, blue, and linking motifs respectively. All the sticky-end sequences and the fluorescent modifications are indicated in the color of the population to which the sequence belongs. The two spacer bases (TT, rUrU) are colored in gray. RNA and photolabile sequences are highlighted in orange and magenta, respectively. Ribonucleotides are preceded with a letter "r". Positions of nitrobenzyle linkers are marked with an asterisk. The free adenine base is highlighted in bold.

| Name | Sequence(5'-3') |
|---|---|
| G-1 | GCTCGAGCCAGTGAGGACGGAAGTTTGTCGTAGCATCGCACC |
| G-2 | GCTCGAGCCAACCACGCCTGTCCATTACTTCCGTCCTCACTG |
| G-3 | GCTCGAGCGGTGCGATGCTACGACTTTGGACAGGCGTGGTTG |
| B-1 | CTCGCGAGAAAGGAACTCTCCGCGTTGACAAAGCCGACACGT |
| B-2 | CTCGCGAGGCCTCTGTGTCGCATCTTCGCGGAGAGTTCCTTT |
| B-3 | CTCGCGAGACGTGTCGGCTTTGTCTTGATGCGACACAGAGGC |
| L-2 | CTCGCGAGCTCAGAGAGGTGACAGCATTCCGTTCCGTTAGTCCAGC |
| L-3 | CTCGCGAGCCATGGTCCCAAGTGATGTTTGCTGTCACCTCTCTGAG |
| L-5 | GCTCGAGCCAGACGTCACTCTCCAACTTCGCAAATTTACAGCGCCG |
| L-6 | GCTCGAGCGTGCTGGCATACCTGACTTTGTTGGAGAGTGACGTCTG |
| L-1-RNA | CTCGCGAGGCTGGACTAACGGArArCrGrGrUrUrArGrUrCAGGTATGCCAGCAC |
| L-4-RNA | GCTCGAGCCGGCGCTGTAAATTrUrGrCrGrUrUrCrArUrCACTTGGGACCATGG |
| L-1-Photo | CTCGCGAGGCTGGACTAACGGA*ACGGT*TAGTC*AGGTATGCCAGCAC |
| L-4-Photo | GCTCGAGCCGGCGCTGTAAATT*TGCGT*TCATC*ACTTGGGACCATGG |
| G-free-1 | GCTCGAGC**A**CAGTGAGGACGGAAGTTTGTCGTAGCATCGCACC |
| G-free-2 | GCTCGAGC**A**CAACCACGCCTGTCCATTACTTCCGTCCTCACTG |
| G-free-3 | GCTCGAGC**A**GGTGCGATGCTACGACTTTGGACAGGCGTGGTTG |
| B-free-1 | CTCGCGAG**A**AAAGGAACTCTCCGCGTTGACAAAGCCGACACGT |
| B-free-2 | CTCGCGAG**A**GCCTCTGTGTCGCATCTTCGCGGAGAGTTCCTTT |
| B-free-3 | CTCGCGAG**A**ACGTGTCGGCTTTGTCTTGATGCGACACAGAGGC |
| L-2-Cy3 | [Cy3]-CTCAGAGAGGTGACAGCATTCCGTTCCGTTAGTCCAGC |
| L-5-Cy5 | [Cy5]-CAGACGTCACTCTCCAACTTCGCAAATTTACAGCGCCG |
| L-2-ATTO647N | [ATTO647N]-CTCAGAGAGGTGACAGCATTCCGTTCCGTTAGTCCAGC |
| L-5-ATTO488 | [ATTO488]-CAGACGTCACTCTCCAACTTCGCAAATTTACAGCGCCG |
| L-2-ATTO488 | [ATTO488]-CTCAGAGAGGTGACAGCATTCCGTTCCGTTAGTCCAGC |
| L-5-ATTO647N | [ATTO647N]-CAGACGTCACTCTCCAACTTCGCAAATTTACAGCGCCG |
| G-2-FAM | [6-FAM]-CAACCACGCCTGTCCATTACTTCCGTCCTCACTG |
| B-2-DY405 | [DY405]-GCCTCTGTGTCGCATCTTCGCGGAGAGTTCCTTT |

### 3 Table S2: DNA mixing ratios

Table S2: DNA mixing ratios.
The standard mix for our DNA droplets is shown below. Depending on the DNA droplet type, the mixture can be adjusted. For example: photocleavable droplets consist of L-1-Photo and L-4-Photo while enzyme-labile droplets contain L-1-RNA and L-4-RNA. For different combinations shown in Figure 4, all three strands of G- or B- will be replaced with G-free- and B-free-strands.
For labelling, 30% of the non-fluorescent strands will be replaced with the mentioned fluorescent strands.

| Strand name | Final concentration [µM] |
|---|---|
| G-1 | 5 |
| G-2 | 5 |
| G-3 | 5 |
| B-1 | 5 |
| B-2 | 5 |
| B-3 | 5 |
| L-1 | 1.65 |
| L-2 | 1.15 |
| L-3 | 1.65 |
| L-4 | 1.65 |
| L-5 | 1.15 |
| L-6 | 1.65 |
| L-2-ATTO647N | 0.5 |
| L-5-ATTO488 | 0.5 |

### 4 Figure S1: Enzyme-induced segregation of DNA droplets in water-in-oil droplets

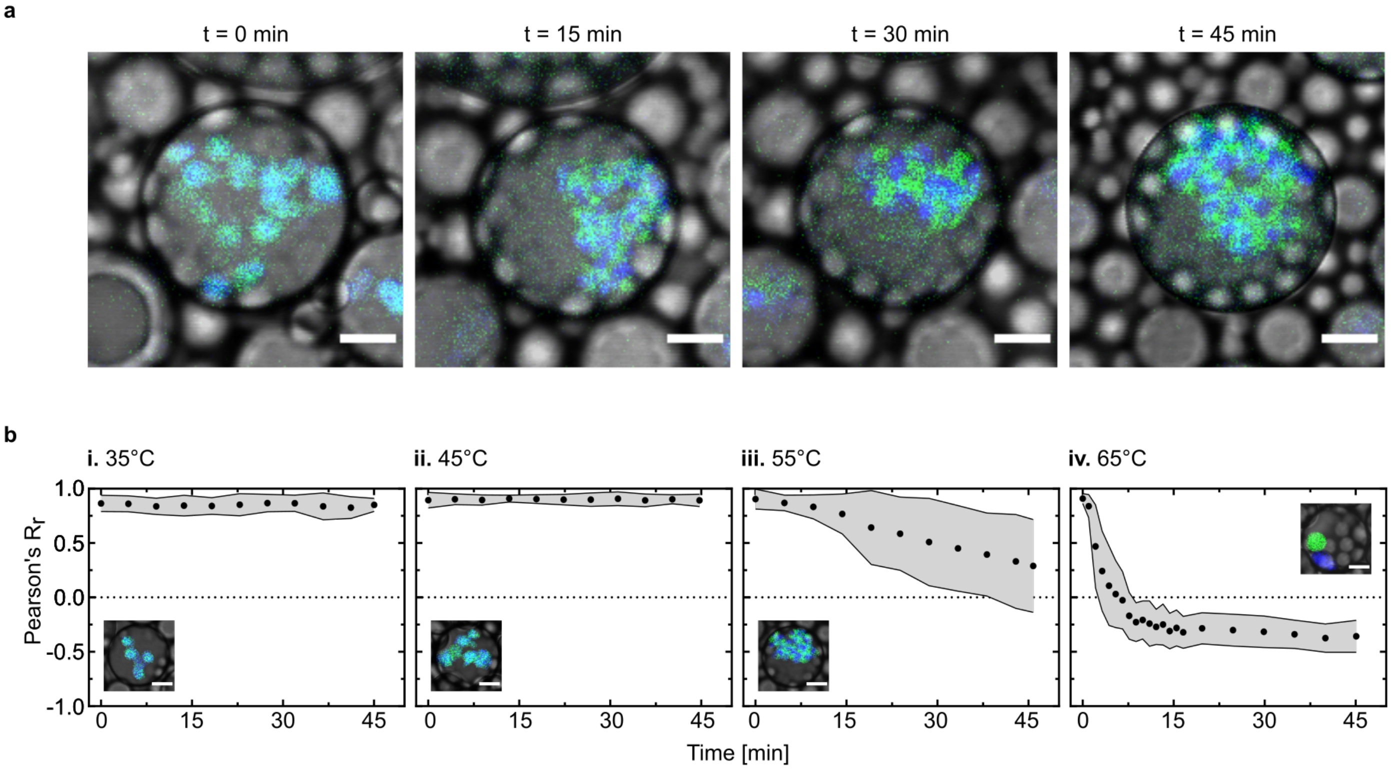


Figure S1: Enzyme-induced segregation of DNA droplets in water-in-oil droplets. a) Time series of DNA droplet segregation triggered by enzymatic activity of RNase A in a water-in-oil droplet (fluorescence and brightfield overlay). The two DNA populations were labeled with ATTO-488 ($\lambda_{ex} = 488\,\mathrm{nm}$) and ATTO-647N ($\lambda_{ex} = 640\,\mathrm{nm}$). b) The enzymatic segregation process is observed over time in water-in-oil droplets at different temperatures (**i-iv**) ($t = 45\,\mathrm{min}$, $T = 35 - 65\,^\circ\mathrm{C}$, $10\,^\circ\mathrm{C}$ step) and colocalization values (Pearson's $R_r$) are plotted (mean ± s.d, $n > 60$ water-in-oil droplets). Insets: Overlay of confocal fluorescence and brightfield images of representative water-in-oil droplets in each condition. The fluorescent pixels outside of the DNA droplets are removed for visibility. Scale bars: 10 µm.

### 5 Figure S2: Simulated DNA droplet segregation in confinement

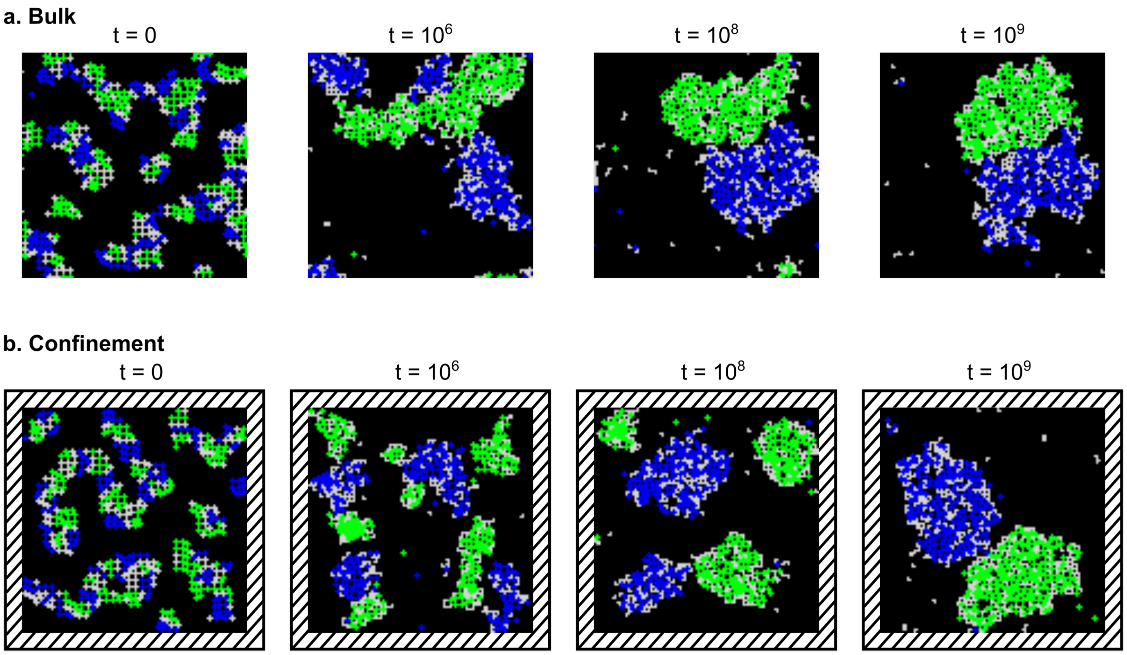


Figure S2: Original images of Figure 2c. Snapshots of DNA droplets undergoing segregation with periodic boundary conditions, mimicking experimental bulk condition (a) and within rigid boundary, mimicking experimental confinement (b). The gray particles are set in its original color.

#### 6 Figure S3: DNA droplet segregation after a temperature rise to 55 °C

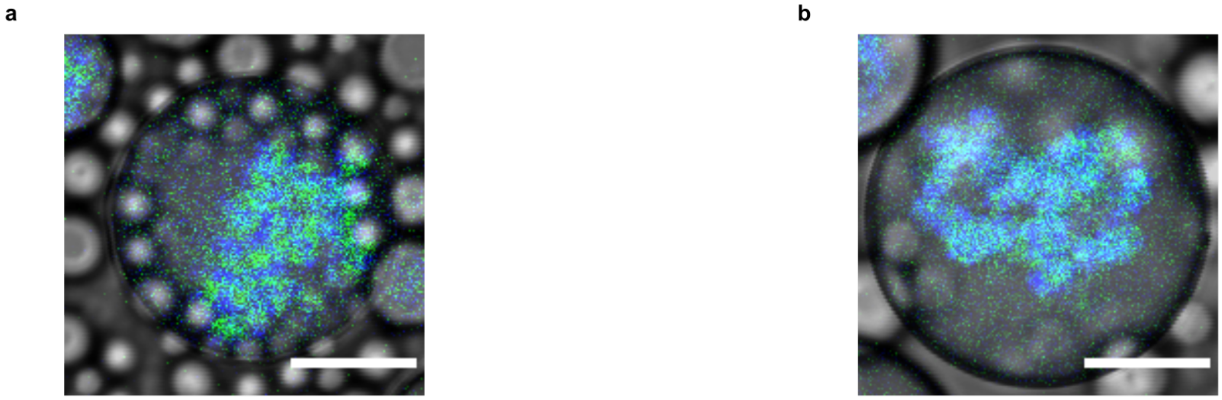


Figure S3: DNA droplet segregation after a temperature rise to 55 °C. After photocleavage and incubation for 45 min at either 35 °C (a) or 45 °C (b), the temperature was increased to 55 °C for either 5 min (a) or 7 min (b). Confocal overlay of brightfield and fluorescence channels of a representative water-in-oil droplet for each condition is shown. Scale bars: 20 µm

### 7 Figure S4: Time series of DNA droplet segregation during incubation at 65 °C

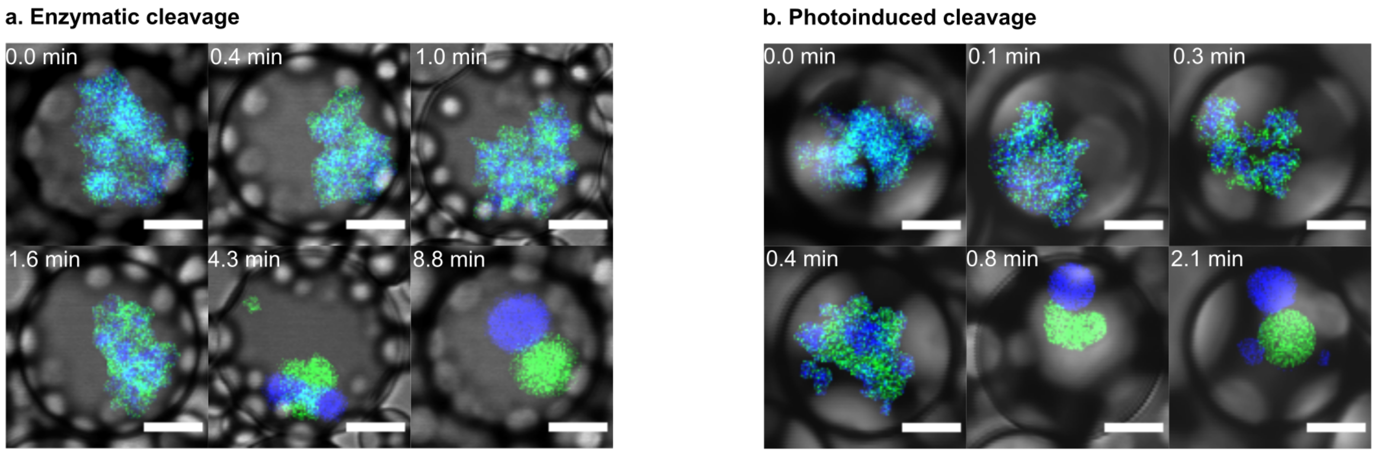


Figure S4: Time series of DNA droplet segregation at 65 °C. Representative confocal microscopy images of different time points showing DNA droplet segregation in water-in-oil droplets (fluorescence and brightfield overlay) after enzyme-induced (a) and photoinduced (b) cleavage. The fluorescent pixels outside of the DNA droplets were removed for visibility. The two DNA populations were labeled with ATTO-488 ($\lambda_{ex}$ = 488 nm) and ATTO-647N ($\lambda_{ex}$ = 640 nm). Scale bars: 10 µm.

### 8 Figure S5: Comparison of DNA droplet segregation process at 55 °C and 65 °C

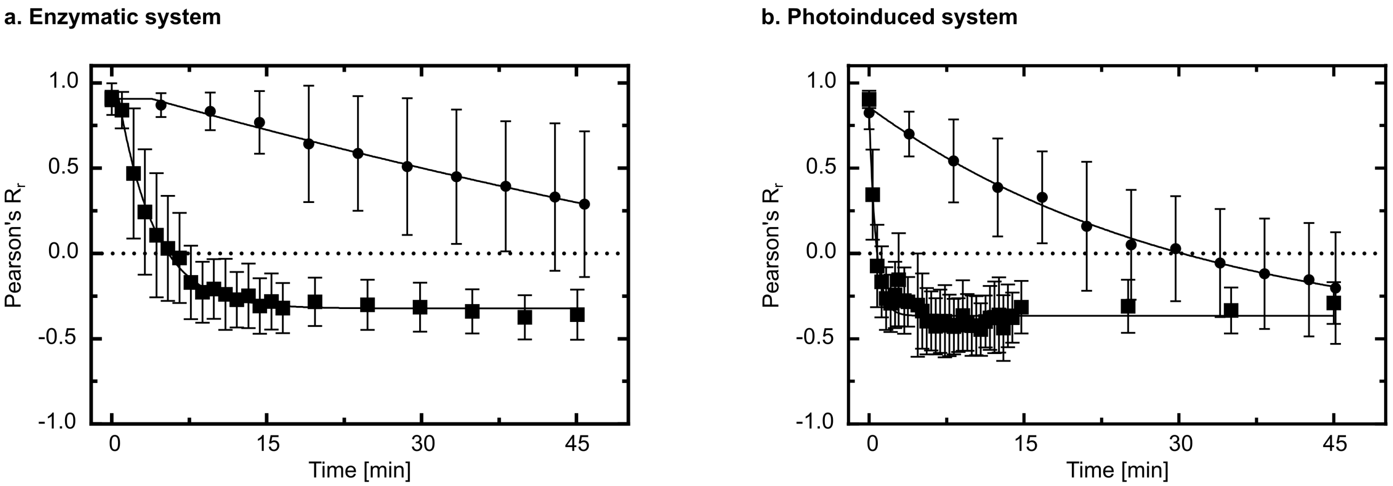


Figure S5: Comparison of DNA droplet segregation process at 55 °C and 65 °C. The data has been extracted from Figure 3a and Figure S1b. Pearson's $R_r$ values over time for segregation process at 55 °C (circles) and 65 °C (squares) are fitted by a plateau followed by a one phase decay equation for enzymatic system (a) and by a one phase decay equation for photoinduced system (b). An extra sum-of-squares F test was performed to compare the fitted curves for data collected at 55 °C and at 65 °C (Y0 is shared value for all data sets, K > 0). P values are < 0.0001 for both systems.

### 9 Figure S6: Overview of DNA droplet segregation after incubation at 55 °C and 65 °C

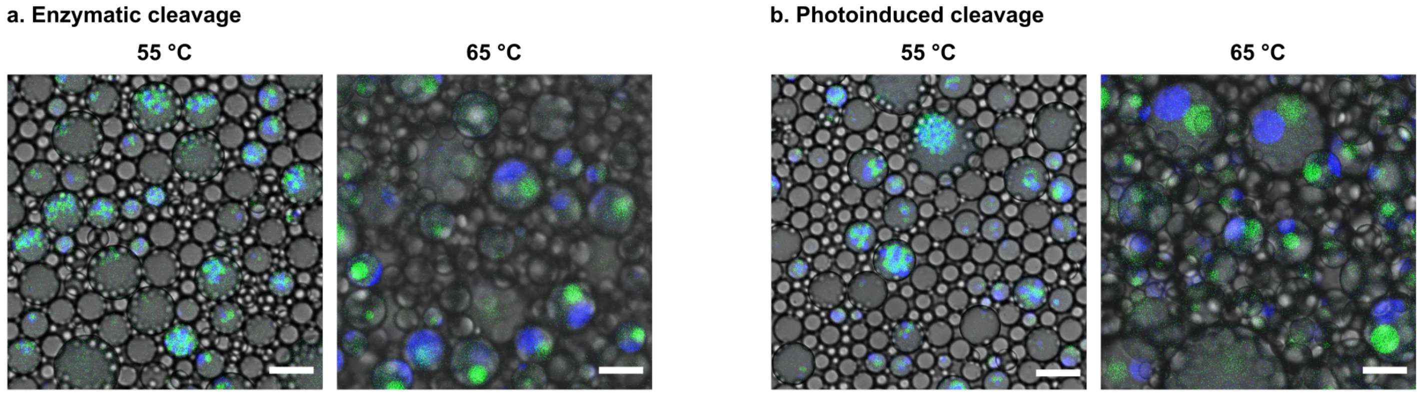


Figure S6: Overview of DNA droplet segregation after incubation at 55 °C and 65 °C. Overlay of brightfield and fluorescence confocal images showing DNA droplet segregation in multiple water-in-oil droplets in the same field of view. DNA droplets are cleaved using RNase A (a) or light illumination (b) and incubated for 45 min at 55 °C or 65 °C. The two DNA populations were labeled with ATTO-488 ($\lambda_{ex} = 488$ nm) and ATTO-647N ($\lambda_{ex} = 640$ nm). Scale bars: 20 µm.

### 10 Figure S7: Melting profile of the sticky end sequences of the two DNA populations

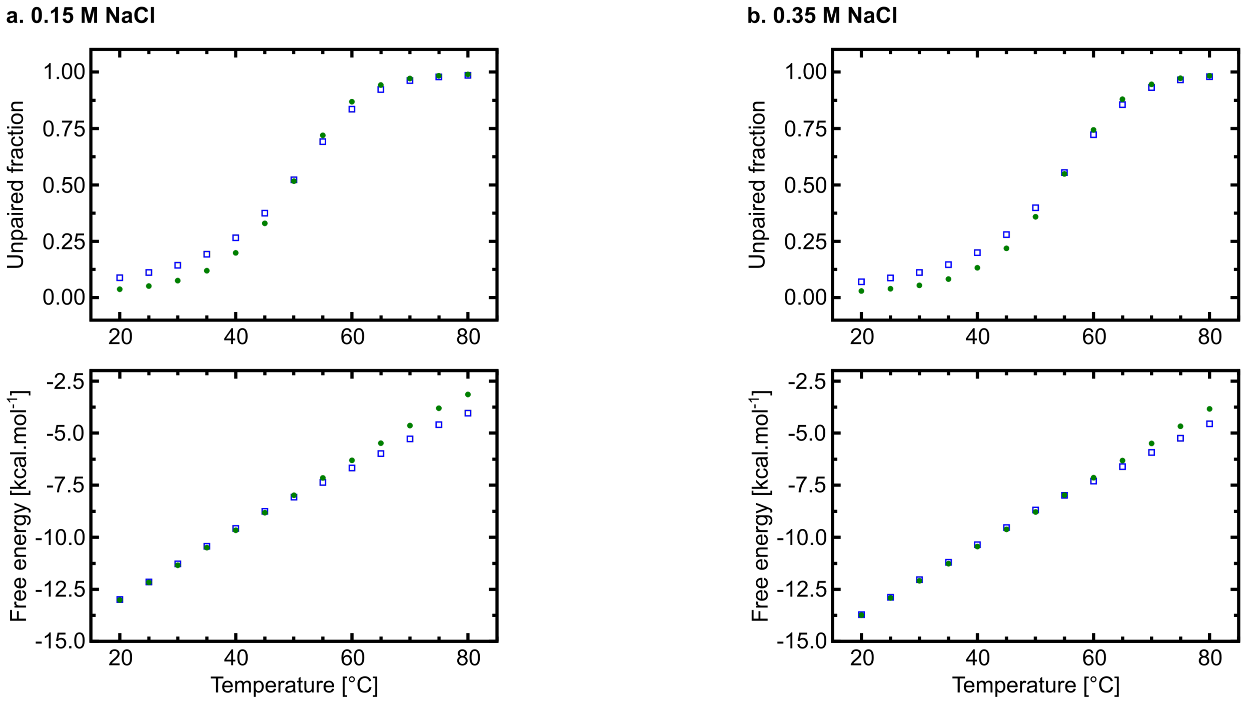


Figure S7: Melting profile of the sticky end sequences of the two DNA populations. The unpaired fraction and free energy of base pairing are plotted over different temperatures (20 −80 °C, 5 °C step) for the sticky end sequences of the green population (show in green circles) and the blue population (shown in blue empty squares). The melting profiles were simulated using NUPACK [2] for 0.15 M NaCl (a) and 0.35 M NaCl (b) with a strand concentration of 160 µM.

#### 11 Figure S8: Effects of the addition of one free nucleobase on droplet formation from each population

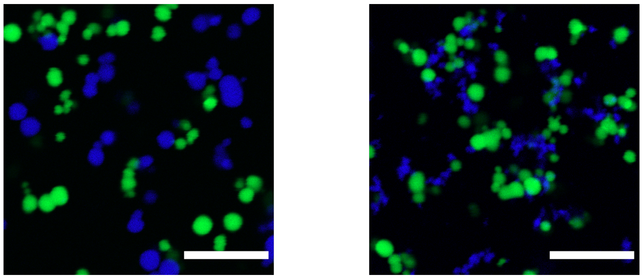

Figure S8: Introduction of one free nucleobase hinders DNA droplet formation of the blue population. Overlay of confocal images of the DNA droplets. The two populations were formed together without the linking motifs (see Experimental Section, Supporting Information) using the original strands (left) or the modified strands (right) where one free base is inserted between the sticky end and the Y-motif arm sequence. The two DNA populations were labeled with FAM ($\lambda_{ex} = 488\,\text{nm}$) and DY405 ($\lambda_{ex} = 405\,\text{nm}$). Scale bars: 20 µm.

#### 12 Figure S9: Effect of the fluorophores on DNA droplet stability

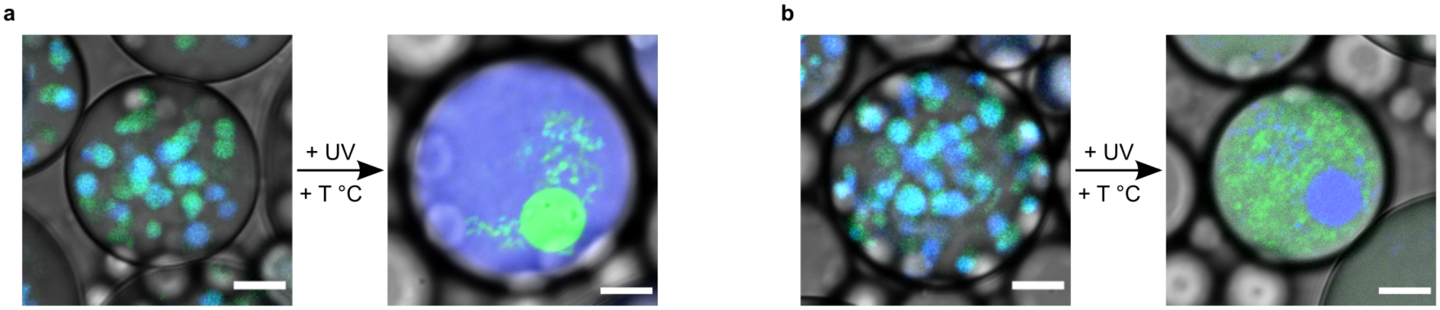


Figure S9: DNA droplet stability after exchanging fluorophores between the two populations. Confocal overlay of brightfield and confocal images showing DNA droplet stability inside water-in-oil droplets before (left) and after cleavage and incubation at 55 °C for 105 min (right). The green and blue populations were labeled with ATTO-488 ($\lambda_{ex} = 488$ nm) and ATTO-647N ($\lambda_{ex} = 640$ nm) respectively (a) and vice versa (b). The color of the two populations are switched in (b) to highlight that DNA droplet stability is independent of the fluorophore choices. Scale bars: 10 µm.

#### 13 Figure S10: Effects of the addition of one free nucleobase on DNA droplet formation

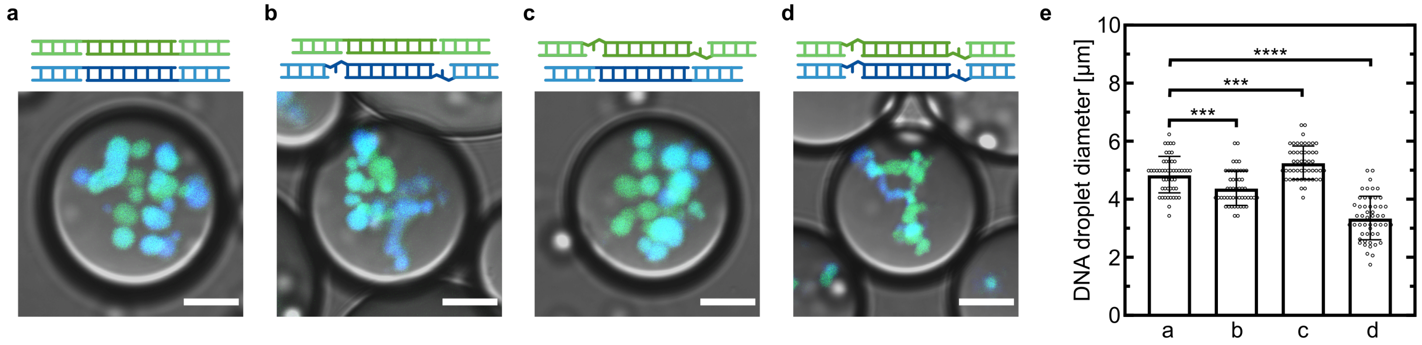


Figure S10: Introduction of one free nucleobase alters the DNA droplet formation process. Overlay of brightfield and fluorescence images of encapsulated DNA droplets in water-in-oil droplets. DNA droplet formation from the four combinations mentioned in Figure 4a was observed (a-d). The droplet diameter of each combination was analyzed (e), showing significant changes in droplet size among the conditions (t-test p-value: p-value $< 0.001$ (***), p-value $< 0.0001$ (****)). The two DNA populations were labeled with ATTO-488 ($\lambda_{ex} = 488\,\mathrm{nm}$) and ATTO-647N ($\lambda_{ex} = 640\,\mathrm{nm}$). Scale bars: 10 µm.

#### 14 Figure S11: Effects of the addition of one free nucleobase on DNA droplet integrity after light cleavage

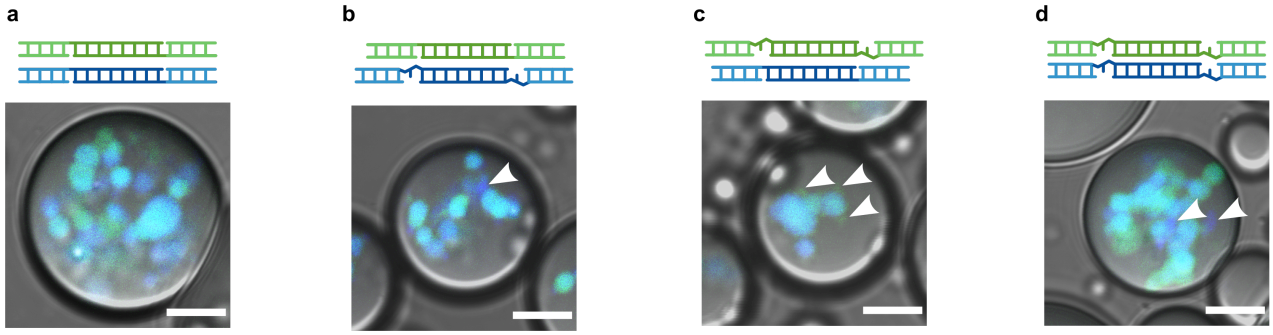


Figure S11: Addition of one free nucleobase introduces aggregations of the modified population after photocleavage. Overlay of brightfield and fluorescence images of the encapsulated DNA droplets in water-in-oil droplets after photocleavage. The DNA droplets were made from different strand combinations as desciribed in Figure 4a (i-iv). The aggregations are highlighted with white arrowheads. In the case where both sticky end sequences were modified, only aggregations of the blue population were observed. The two DNA populations were labeled with ATTO-488 ($\lambda_{ex}$ = 488 nm) and ATTO-647N ($\lambda_{ex}$ = 640 nm). Scale bars: 10 µm.

#### 15 Figure S12: DNA droplet segregation in GUVs triggered by RNase A activity

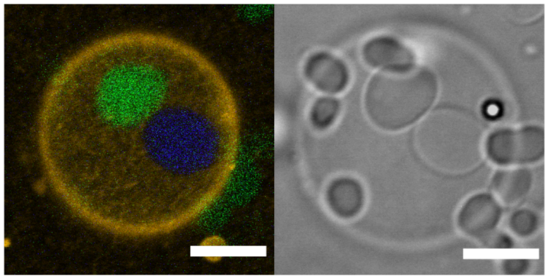

Figure S12: DNA droplet segregation triggered by enzymatic activity in GUVs. Fluorescence overlay (left) and brightfield (right) images showing segregated DNA droplet inside GUV after incubation at 50 °C for 30 min. The green population was labeled with FAM ($\lambda_{ex} = 488$ nm) and Cy5 ($\lambda_{ex} = 651$ nm). The blue population was labeled with DY405 ($\lambda_{ex} = 405$ nm). The lipid membrane was labeled with Liss Rhod PE ($\lambda_{ex} = 561$ nm). Scale bars: 10 µm.

### 16 Figure S13: Local light-induced DNA droplet segregation in GUVs

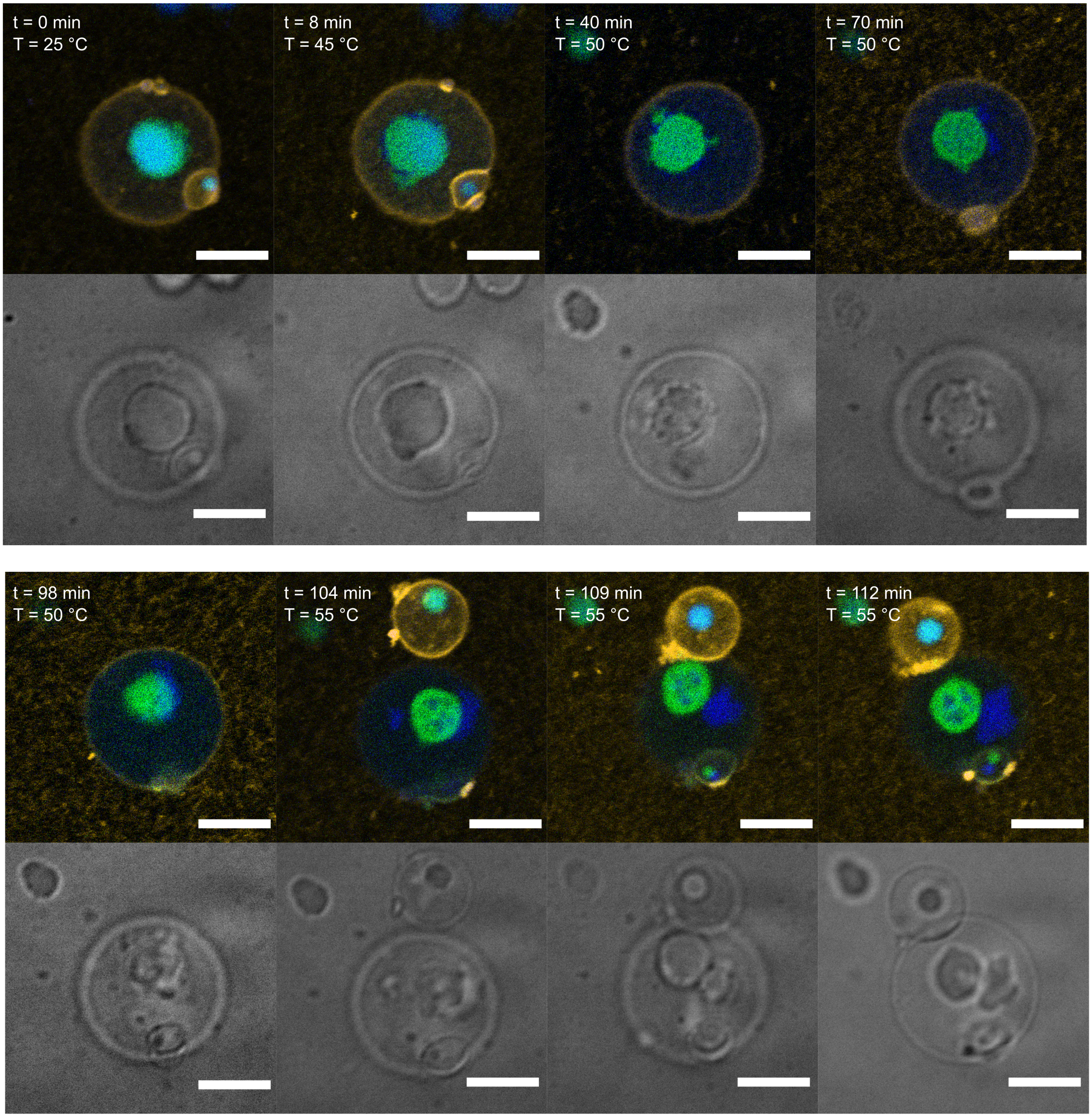


Figure S13: Time series of DNA droplet segregation locally triggered by light in GUVs. Fluorescence overlay (upper panel) and brightfield (lower panel) images showing DNA droplet segregation process inside GUV after photocleavage ($t = 1$ min, $\lambda = 405$ nm, 5 mW power). From 100 min, an unilluminated GUV entered the field of view (see also Video S4). The absence of DNA droplet segregation the unilluminated GUV highlights the spatiotemporal control over the cleavage. The two DNA populations were labeled with ATTO-488 ($\lambda_{ex} = 488$ nm) and ATTO-647N ($\lambda_{ex} = 640$ nm). The lipid membrane was labeled with Liss Rhod PE ($\lambda_{ex} = 561$ nm). Scale bars: 10 µm.

#### 17 Supporting videos

##### 17.1 Video S1: Photoinduced DNA droplet segregation in bulk at 55 °C

Confocal time lapse of DNA droplet segregation in bulk at 55 °C. The droplets were illuminated using an UV lamp for 5 min to induce photocleavage (see Experimental Section). The fluorescent and brightfield channels were split and are presented in separated panels (upper left: green population, upper right: blue population, lower right: brightfield channel showing both populations). The composite of the three channels are shown in the lower left panel. The two DNA populations were labeled with ATTO-488 ($\lambda_{ex}$ = 488 nm) and ATTO-647N ($\lambda_{ex}$ = 640 nm).

##### 17.2 Video S2: Photoinduced DNA droplet segregation in the confinement of water-in-oil droplets at 55 °C

Confocal time lapse of DNA droplet segregation in water-in-oil droplets at 55 °C. The droplets were illuminated using an UV lamp for 5 min to induce photocleavage (see Experimental Section). Images are composite of fluorescence and brightfield channels. The two DNA populations were labeled with ATTO-488 ($\lambda_{ex}$ = 488 nm) and ATTO-647N ($\lambda_{ex}$ = 640 nm).

##### 17.3 Video S3: Local light-triggered DNA droplet segregation in GUV

Confocal time lapse of the full segregation of a DNA droplet inside GUV. A single DNA droplet inside GUV fully segregated into two daughter DNA droplets upon illumination ($\lambda$ = 405 nm, $t$ = 30 s, $I$ = 5 mW) using the temperature profile shown in Figure 4c. The fluorescent and brightfield channels were split and are presented in separated panels (upper left: green population, upper right: blue population, lower right: brightfield channel showing both DNA populations and the lipid membrane). The composite of the two fluorescence channels is shown in the lower left panel. The two DNA populations were labeled with ATTO-488 ($\lambda_{ex}$ = 488 nm) and ATTO-647N ($\lambda_{ex}$ = 640 nm).

##### 17.4 Video S4: Local light-triggered DNA droplet segregation in fluorescently labelled GUV

Confocal time lapse of DNA droplet segregation inside a fluorescently labelled GUV. DNA droplet segregation inside GUV was triggered upon illumination ($\lambda$ = 405 nm, $t$ = 1 min, $I$ = 5 mW) and heat application. The applied temperature profile is as follows: 1: temperature increase from 25 °C to 45 °C at an increment of 0.1 °C s$^{-1}$; 2: incubate at 45 °C for 5 min; 3: temperature increase from 45 °C to 50 °C at an increment of 0.1 °C s$^{-1}$; 4: incubate at 50 °C for 90 min; 5: temperature increase from 50 °C to 55 °C at an increment of 0.1 °C s$^{-1}$; 6: incubate at 55 °C until the end of the experiment. Composites of fluorescent channels are shown: lipid membrane and green population (upper left), lipid membrane and blue population (upper right), lipid membrane and the two DNA populations (lower left). The brightfield channel is shown in the lower right. The two DNA populations were labeled with ATTO-488 ($\lambda_{ex}$ = 488 nm) and ATTO-647N ($\lambda_{ex}$ = 640 nm). The lipid membrane was labeled with Liss Rhod PE ($\lambda_{ex}$ = 561 nm).
